# The Barcode, UMI, Set format and BUStools

**DOI:** 10.1101/472571

**Authors:** Páll Melsted, Vasilis Ntranos, Lior Pachter

## Abstract

We introduce the **B**arcode-**U**MI-**S**et format (BUS) for representing pseudoalignments of reads from single-cell RNA-seq experiments. The format can be used with all single-cell RNA-seq technologies, and we show that BUS files can be efficiently generated. BUStools is a suite of tools for working with BUS files and facilitates rapid quantification and analysis of single-cell RNA-seq data. The BUS format therefore makes possible the development of modular, technology-specific, and robust workflows for single-cell RNA-seq analysis.

## Introduction

The analysis of single-cell RNA-seq data begins with three related computational tasks: read assignment to transcripts, cell determination, and molecule identification [1,2]. These tasks are accomplished by utilizing information encoded in reads produced from single-cell RNA-seq experiments. While the exact nature of the encoded information is technology specific, the components are universal: “cell barcodes” are short sequences that identify the cells of origin for each read, “unique molecular identifiers” (UMIs) are sequences that identify the molecule of origin for each read, and finally transcripts of origin are encoded via reverse transcribed cDNA sequences. Read assignment is typically accomplished by alignment of reads to a reference genome or transcriptome. Cell determination, which is the process of determining the valid cells in an experiment along with the reads associated to those cells, involves grouping of similar barcodes that appear to differ only due to sequencing or synthesis error, collation of the reads associated with those barcodes, and determination of valid cells according to alignment statistics of the reads associated to them. Molecule identification, which is the process of determining which reads originated from the same molecule, consists of collapsing read counts when UMI sequences match in reads that have been assigned to a single transcript or gene from one cell. The challenges that must be overcome to efficiently and accurately solve the assignment, determination and identification problems are both computational and algorithmic. On the one hand, increasing throughput of experiments [3] has resulted in large numbers of reads that make it difficult to perform tasks such as comparison of barcodes or alignment of reads. At the same time, problems of cell determination and molecule identification can benefit from innovative algorithmic ideas, e.g. [4]. Current commonly used single-cell RNA-seq workflows [5–8] confound these challenges and this has led to numerous drawbacks: software packages are frequently technology specific, the replacement of individual steps when better methods become available can be difficult, and running time/memory requirements may limit the scale of experiments that can be analyzed.

We introduce a new file format for representing single-cell RNA-seq data called BUS that is an abbreviation for Barcode, UMI, and Set. BUS format consists of a binary representation of barcode and UMI sequences from single-cell RNA-seq reads, along with sets of equivalence classes of transcripts [9] obtained by pseudoalignment of the reads to a reference transcriptome [10]. BUS files can be rapidly and efficiently produced, and we provide a solution based on kallisto (pachterlab.github.io/kallisto) that can create BUS files using data from any of seven different single-cell RNA-seq technologies. However, the BUS format is neither technology nor software dependent. The utility of BUS files lies in their compact representation of the key information from single-cell RNA-seq experiments that is needed for quantification. They enable a modular approach to single-cell RNA-seq processing that separates compute intensive read assignment from algorithmically demanding cell determination and molecule identification. Importantly, BUS also decouples technology dependencies (Figure 1a). By virtue of avoiding the explicit representation of transcriptome sequences, BUS is also useful for sharing the important aspects of raw data in a way that removes identifying genotypes.

## Methods

### The BUS format

The BUS format consists of two files: the first describes the mapping of equivalence classes to sets to transcripts and the second records the BUS tuples in binary format. Each record of the BUS file consists of a barcode and UMI sequence encoded using a 2-bit format, the equivalence class, count and optional flags (Figure 1b). There is an inherent limit on the size of the barcode and UMI sequences set at 32bp each. Each sequenced fragment corresponds to a single BUS record and no information about read names is stored.

### kallisto bus

A BUS file can be produced by any alignment or pseudoalignment method. As a proof of principle and to illustrate the versatility and utility of BUS, we implemented a bus command for the kallisto pseudoalignment software [9] that will output BUS format from single-cell RNA-seq data. kallisto bus can accept as input single-cell RNA-seq generated from 10x v1 [11], 10x v2 [11], 10x v3 [11], inDrops [7], Drop-seq [8], CEL-seq [12], CEL-seq2 [13] SCRB-seq [14] and SureCell [15]; other technology formats can be readily processed by setting options.

### BUStools

BUStools are a software suite developed to manipulate and organize BUS files (https://github.com/BUStools). The tools currently consist of programs for sorting BUS files, merging BUS files, and converting BUS files to a textual representation.

## Results

To demonstrate the utility of BUS we processed 381,992,071 single-cell RNA-seq reads from a 1:1 mixture of fresh frozen human cells (HEK293T) and mouse cells (NIH3T3) produced with 10x technology [4] and hosted on the 10x Genomics website [16]. kallisto bus is more than 50 times faster than CellRanger, processing the dataset in 984 seconds vs. 55,745 seconds with CellRanger, and almost 4 times faster than the 3,786 seconds required with Alevin [6] (Figure 1c). Crucially, the memory requirements of kallisto bus are constant in the number of reads, a feature that reduces cost for processing single-cell RNA-seq data with cloud infrastructure. Furthermore, kallisto bus is sufficiently fast with only 4 threads to provide users with limited compute resources the ability to rapidly process standard datasets in real time.

To illustrate the possibilities created by the modular BUS format, we developed a simple “naïve” post-processing notebook (https://github.com/BUStools). The notebook provides statistics on barcodes, UMIs and gene counts, and can be used for a very rapid assessment of data. Figure 1d shows an example figure from the notebook, namely the distribution of genes detected for the 10x human-mouse 6k dataset.

## Acknowledgments

We thank Fan Gao for helping with the benchmarking of kallisto bus and BUStools. We thank Valentine Svensson for helpful discussions and suggestions, and for compiling a comprehensive reference of single-cell RNA-seq read encodings [17] that we relied on. Jase Gehring, Lynn Yi and Tina Wang provided valuable feedback on an initial kallisto-based single-cell RNA-seq workflow, which motivated the development of the BUS format.

---

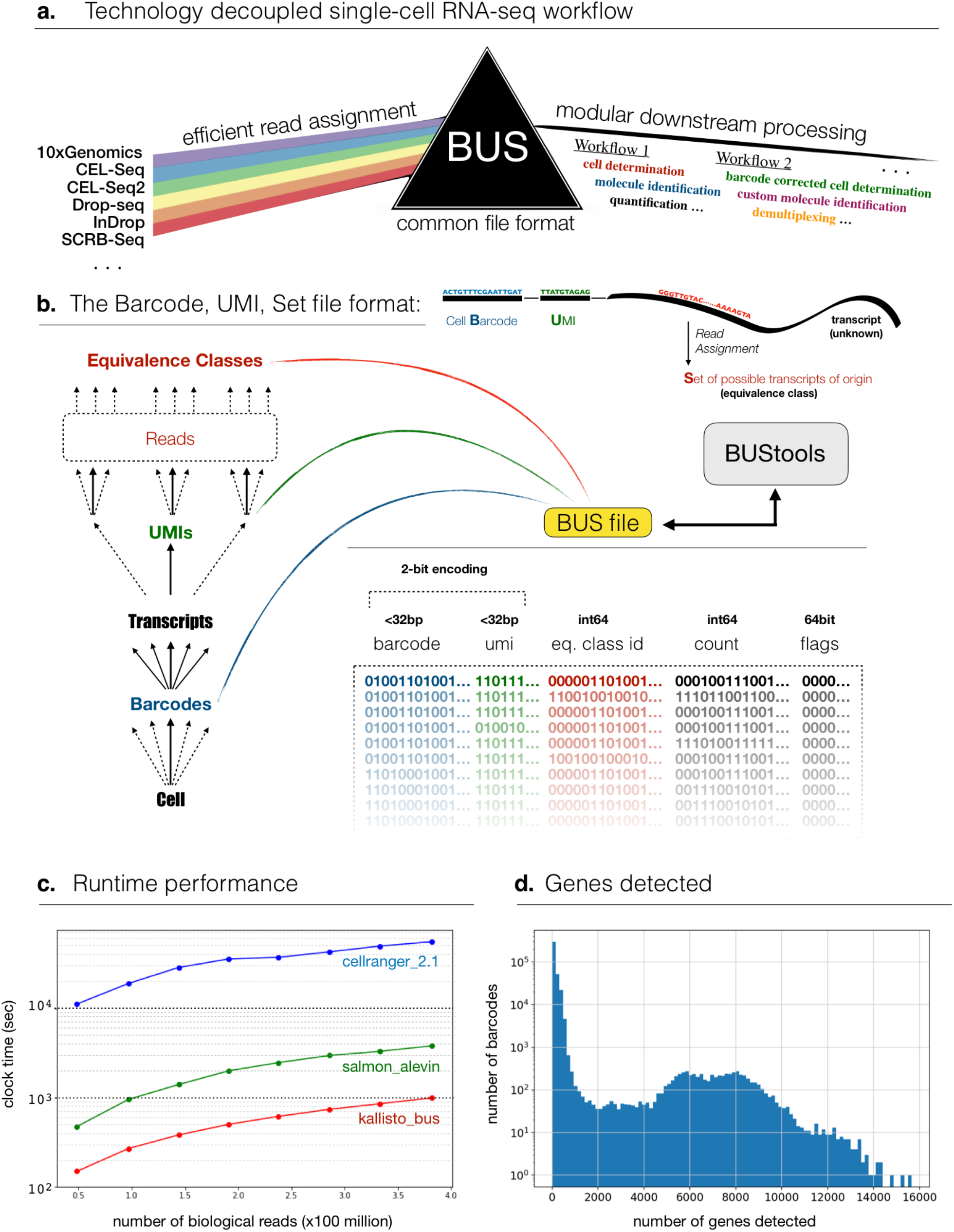

